# Combining Selenocystine Based Fluorescence Assay with Structural Prediction for Enhanced Detection and Analysis of Cystine Stone Pathology

**DOI:** 10.1101/2025.08.29.673186

**Authors:** Xiaobai He, Xinyi Qian, Xiaoguang Zheng, Hong Zhang, Jinbang Shao, Xiaopan Chen, Ruan Qi, Jianxin Lyu, Leixiang Yang, Linjie Chen

**Author notes:** **Corresponding author: Linjie Chen**, PhD, School of Laboratory Medicine and Bioengineering, Hangzhou Medical College, No.8, Yikang ST, Lin’an District, Hangzhou, Zhejiang, 311399, China.; **Leixiang Yang**, PhD, Department of Genetic and Genomic Medicine, Zhejiang Provincial People’s Hospital, Affiliated People’s Hospital, No. 158, Shangtang Road, Gongshu District, Hangzhou, Zhejiang, 310014, China.; **Jianxin Lyu**, School of Laboratory Medicine and Bioengineering, Hangzhou Medical College, No.8, Yikang ST, Lin’an District, Hangzhou, Zhejiang, 311399, China.

## Abstract

**Background:** Cystine stones, a rare but recurrent type of kidney stones, primarily result from cystinuria, an inherited disorder caused by mutations in the genes *SLC3A1* and *SLC7A9*, which encode renal cystine transporters. These mutations impair cystine reabsorption, resulting in elevated urinary cystine concentrations and stone formation. Current diagnostic methods are limited, particularly for detecting molecular-level dysfunctions. We aimed to develop a nonradioactive method for functional assessment of cystine transporters and mutation-specific pathologies.

**Methods:** We designed an innovative diagnostic approach combining a selenocystine-based fluorescence assay with structural predictions using AlphaFold to rapidly and accurately assess cystine transporter function and the molecular impacts of genetic mutations.

**Results:** Our assay demonstrated comparable transport efficiencies of cystine and selenocystine by the SLC3A1/SLC7A9 complex, and effectively differentiated mild, moderate, and severe functional impairments associated with known clinical mutations, including A354T and P482L. Structural modeling further provided mechanistic insights into mutation-induced dysfunctions.

**Conclusion:** This integrated approach—combining a sensitive selenocystine fluorescence assay with AI-powered structural analysis—enables rapid, precise diagnosis of cystinuria variants and delivers mechanistic insights for personalized therapeutic strategies.

## 1. Introduction

Cystine stones, account for 1-2% of adult kidney stones and up to 10% in pediatric patients, resulting from cystinuria, an autosomal recessive disorder caused by mutations in SLC3A1 and SLC7A9 genes. These mutations impair renal cystine reabsorption, promoting crystallization (1–4). Early diagnosis and targeted intervention are critical to prevent the progression of stone formation and kidney damage. However, current diagnostics remains limited, particularly when it comes to detecting cystine transport dysfunctions at the molecular level. Functional analysis of mutations by uptake measurements with radioisotope labeled amino acids, such as [35S]-L-cystine(5), [14C]-L-cystine(6, 7) significantly limit the screening at large scale level due to the safety risks of radioisotope manipulation.

Advances in molecular structural biology have opened new avenues for understanding the pathophysiology of cystine stones. In particular, research on the SLC3A1/SLC7A9 complex structure has significantly improved our understanding of the molecular mechanisms underlying cystine transport (7–9). Additionally, innovations in protein structure prediction, such as the use of AlphaFold, provide unprecedented opportunities to model the effects of genetic mutations on cystine transporter proteins (10). These structural insights are invaluable for understanding how mutations impact protein function, and by extension, cystine reabsorption in the kidneys.

In this study, we present an innovative approach that combines a fluorescence-based cystine transport assay with AlphaFold-based structure prediction to provide a rapid and detailed molecular pathological analysis of cystine stones. The fluorescence assay offers a sensitive and efficient method for detecting cystine transport activity, while AlphaFold enables us to predict structural alterations in cystine transporter proteins caused by specific mutations. Our approach not only improves upon traditional detection methods but also bridges the gap between genetic mutations and their functional consequences at the protein level. By elucidating the molecular mechanisms underlying cystine stone formation, this study will help to pave the way for developing more targeted diagnostic and therapeutic strategies for cystinuria.

## 2. Materials and Methods

### 2.1 Gene Cloning

The cDNA sequences for SLC3A1 (NM_000341.4) and SLC7A9 (NM_001243036.2) were synthesized and cloned into the pLV3-CMV-MCS-3FLAG-EF1alpha-copGFP-PURO plasmid vector. In addition to the wild-type constructs, single point mutation plasmids for SLC7A9 were generated using site-directed mutagenesis. The mutations introduced were A70V, G105R, V170M, A182T, A354T, R333W, P482L and P482G. For SLC3A1, a point mutation (M467T) was also introduced. The wild-type and mutant SLC7A9 and SLC3A1 constructs were confirmed by sequencing.

### 2.2 Cell Lines and transfection

HEK293 cells were cultured in DMEM supplemented with 10% fetal bovine serum (FBS, Sigma, USA), 1% penicillin-streptomycin, and incubated at 37℃ in a humidified incubator with 5% CO_2_. These plasmids were transfected into HEK293 cells using Lipofectamine 2000, following the manufacturer’s protocol. Forty-eight hours post-transfection, cells transiently expressing the exogenous wild-type or mutant transporters were re-seeded into multi-well plates for analysis.

### 2.3 Atomic Absorption Spectroscopy

HEK293 cells transiently co-expressing SLC3A1 and SLC7A9 were washed three times with cystine-free Hanks’ Balanced Salt Solution (HBSS) to deplete endogenous cystine. Cells were then incubated with 200 μM selenocystine in cystine-free HBSS at 37°C for 30 minutes. Following incubation, cells were washed three times with ice-cold HBSS to remove extracellular selenocystine, followed by two washes with ice-cold phosphate-buffered saline (PBS) to minimize salt interference. Washed cells were harvested and digested using microwave-assisted digestion in concentrated nitric acid (HNO_3_) with the following program: 150 ℃ for 10 minutes, followed by 180 ℃ for 20 minutes to mineralize organic material and liberate selenium atoms. The acid was subsequently evaporated to near-dryness under a fume hood. Digested samples were reconstituted in dilute nitric acid (2% v/v) and analyzed by graphite furnace atomic absorption spectroscopy (GFA-6880, Shimadzu Corporation). Selenium content was quantified at its characteristic absorption wavelength (196.0 nm) using a selenium-specific hollow cathode lamp (HAS-1, Guobiao (Beijing) Testing & Certification Co., Ltd). Quantitation was performed against a matrix-matched standard curve generated from certified selenium reference solutions.

### 2.4 Cystine Uptake Assay

HEK293 cells were seeded at a density of 1 × 10⁴ cells per well in a poly-L-lysine coated black, clear-bottom 96-well plate and cultured overnight at 37°C in a humidified 5% CO₂ atmosphere. The following day, the growth medium was removed. Cells were pre-incubated with pre-warmed Hanks’ Balanced Salt Solution (HBSS) at 37°C for 5 minutes to ensure cystine deprivation, then washed twice with cystine-free HBSS. Subsequently, cells were incubated with selenocystine in cystine-free HBSS at 37°C for the indicated time periods; control wells received HBSS without substrate. Following incubation, cells were washed three times with ice-cold PBS and lysed with ice-cold methanol. Cells were then incubated at 37°C for 30 minutes in 100 mM MES buffer (pH 6.0) containing 10 µM Fluorescein O,O’-diacrylate (FOdA, Sigma #7262-39-7) and 200 µM tris (2-carboxyethyl) phosphine hydrochloride (TCEP, Sigma #51805-45-9). The fluorescence intensity was measured using a microplate reader (Infinite M200PRO, TECAN) at an excitation wavelength of 485 nm and an emission wavelength of 535 nm. Background fluorescence, determined from identically treated blank control wells (without substrate), was subtracted from sample well readings.

### 2.5 Structure Prediction of SLC3A1 and SLC7A9

The wild-type structures of SLC7A9 and the SLC3A1-SLC7A9 complex were predicted using the AlphaFold Server platform developed by DeepMind, which provides protein structure prediction services based on the latest version of AlphaFold3, enabling high-speed and high-accuracy predictions without the computational limitations of local hardware. For mutant structure prediction, sequences were then submitted to AlphaFold Server to generate structural predictions for both SLC7A9 mutant monomers and SLC3A1-SLC7A9 heterodimeric mutant complexes. The confidence scores (pLDDT) provided by AlphaFold3 were used to evaluate the reliability of the predicted models. The predicted structures were integrated and analyzed for conformational changes using UCSF ChimeraX (version 1.7.1).

### 2.6 Molecular Docking of Cystine with SLC7A9

The structure file of cystine was downloaded from PubChem (https://pubchem.ncbi.nlm.nih.gov/). The cystine molecule and the AlphaFold3-predicted SLC7A9 protein structure were prepared for docking using AutoDock Tools, including the removal of water molecules and addition of hydrogen atoms. Molecular docking was then performed using AutoDock Vina (version 1.1.2) with the following parameters: the docking boxes were centered on three key regions of SLC7A9 - the putative entry site (near Gly41), exit site (near Ser54), and interior. A total of 1000 docking runs per region were performed for comprehensive exploration of potential binding modes. All docking results were saved in Protein Data Bank (PDB) format for subsequent analysis.

### 2.7 Statistics

Statistical analyses were performed using GraphPad Prism 8.0 (GraphPad Software, San Diego, CA). Data are presented as mean ± standard deviation (SD). Time-dependent responses were analyzed using linear regression, and dose-dependent relationships were modeled using nonlinear regression (Michaelis-Menten or Agonist vs. response). One-way ANOVA was used for analyzing differences between groups. *P* value < 0.05 was considered as statistically significant (marked as *)

## 3. Results

### 3.1 Selenocystine is Transported into Cells by SLC3A1/SLC7A9 Transporters

SLC3A1/SLC7A9 heterodimers are responsible for the reabsorption of cystine and basic amino acids in renal proximal tubules. Based on structural similarity among amino acids and a recently report that selenocystine could be transported by SLC3A2/SLC7A11 complex (11), we hypothesized that selenocystine, a rare but significant amino acid, could also be transported by SLC3A1/SLC7A9. To validate this hypothesis, we employed Atomic Absorption Spectroscopy (AAS) to quantify selenocystine uptake mediated by the SLC3A1/SLC7A9 heterodimer. As shown in Fig. 1A, selenium was detected exclusively in HEK293 cells co-expressing both SLC3A1 and SLC7A9. However, the time-consuming and complex operation limits its high-throughput application. Therefore, we applied a novel method to monitor SLC3A1/SLC7A9 transporting activity. The principle of this method involves collecting cell incorporated selenocystine, reducing it to selenocysteine, and exposing it to a fluorescent dye containing acrylate moieties (11). Selenocysteine reacts with the acrylate moiety, transforming the dye into a fluorescent state with a specific excitation/emission peak (Fig. 1B).

**Figure 1.**
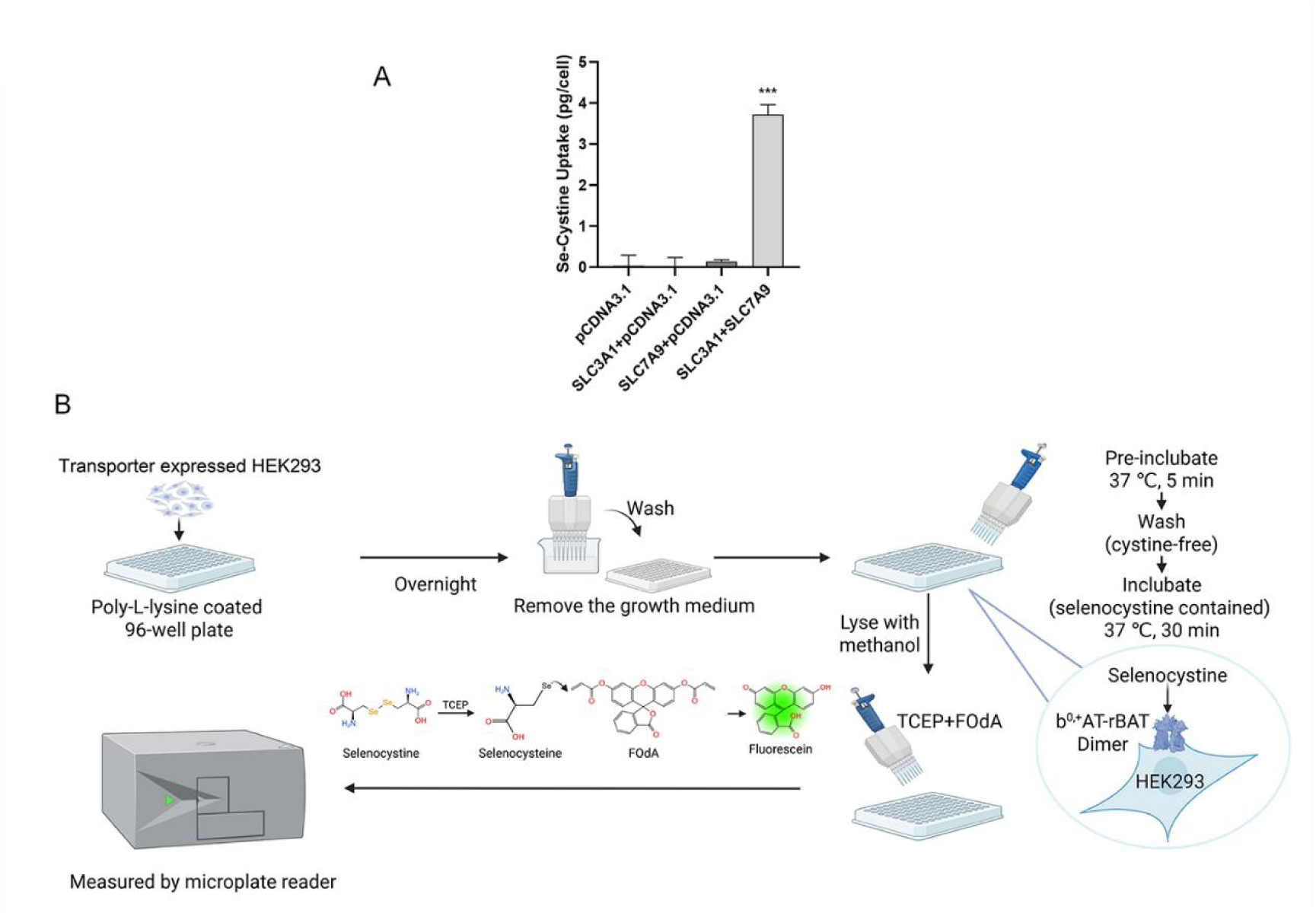
SLC3A1/SLC7A9-dependent selenocystine uptake in HEK293 cells by AAS and fluorescence uptake assay. (A) Cellular selenium content of HEK293 cells was quantified by Atomic Absorption Spectroscopy (AAS) after 30-minute exposure to 200 µM selenocystine. Data represent functional uptake mediated by the SLC3A1/SLC7A9 heterodimer. Methodology details are provided in Materials and Methods. *** *P*<0.001. (B) Graphic Model of Selenocystine-Based Fluorescence Uptake Assay.

Due to the higher reducing activity of the selenol group, selenocystine is significantly more reactive in transforming the dye compared to cystine. As shown in Fig. 2A, the fluorescent signal was more than 100-fold stronger for selenocystine compared to cystine in vitro. We then tested the cellular absorption of selenocystine or cystine in HEK293 cells with SLC3A1/SLC7A9 over-expression. As shown in Fig. 2B, the assay is sensitive enough to detect selenocystine incorporation at a 25 μM concentration incubation, while cystine incorporation is not detectable even at high concentrations (600 μM). Additionally, the selenocystine incorporation was saturable and followed Michaelis-Menten kinetics with a Km of 156.3 μM. Fig. 2C shows that SLC3A1/SLC7A9 transport selenocystine in a time-dependent manner. These results demonstrate the superior sensitivity of this assay for detecting selenocystine transport. To further validate that SLC3A1/SLC7A9 transporters are responsible for selenocystine transport, HEK293 cells were transfected with SLC3A1 or SLC7A9 alone or both. As shown in Fig. 2D, only HEK293 cells co-expressing SLC3A1 and SLC7A9 were able to detect the assay signal.

**Figure 2.**
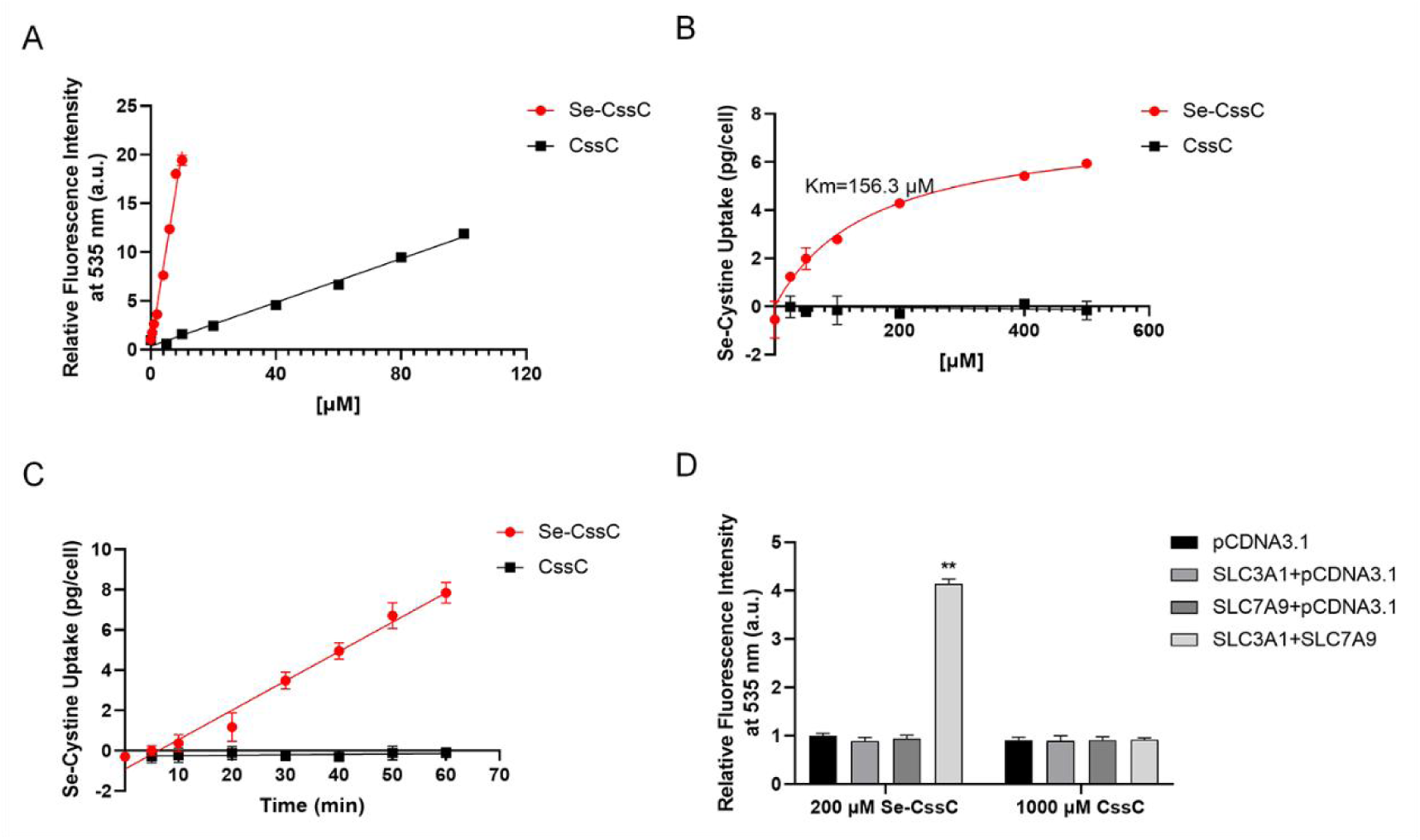
Selenocystine can be uptaken by SLC3A1/SLC7A9 expressing HEK293 cells. (A) Increasing concentration of selenocystine and non-labeled cystine in Hanks’ buffer were reacted with FOdA substrate for 30 min in 37 ℃. Fluorescence signals generated by Selenocystine or non-labeled cystine were measured using a multifunctional microplate reader. (B) Dose-dependent uptake of selenocystine by SLC3A1/SLC7A9-expressing HEK293 cells. Cells were treated with increasing concentrations of selenocystine for 30 minutes, and selenocystine uptake were quantified using a standard curve. Selenocystine transport by SLC3A1/SLC7A9 was fitted to Michaelis-Menten equations, and Km and Vmax were determined by nonlinear regression using Grpahpad Prism software. (C) Time-dependent uptake of selenocystine by SLC3A1/SLC7A9-expressing HEK293 cells. Cells were incubated with 200 μM selenocystine for the indicated durations, and fluorescence intensity was measured to assess uptake over time. (D) HEK293 cells expressing the indicated transporter genes were treated with 200μM selenocystine or 1000 μM cystine for 30 min. Fluorescence intensity was normalized to control conditions. ** *P*<0.01.

### 3.2 Consistent Transporting Characteristics for Selenocystine and Cystine by SLC3A1/SLC7A9

Selenocystine has lower solubility in urine compared to cystine, but its significantly lower concentration in the body prevents crystallization under normal conditions. However, dysfunction of SLC3A1/SLC7A9 transporters leads to impaired cystine reabsorption, resulting in cystine crystallization in the kidneys. To investigate whether SLC3A1/SLC7A9 transporters exhibit similar transport efficiency for cystine and selenocystine, we varied the ratio of cystine to selenocystine in the cell medium. As shown in Fig. 3A, a cystine/selenocystine ratio of 1:1 resulted in a fluorescent signal that was 50% of the signal observed for selenocystine alone. Similarly, a ratio of 3:1 reduced the signal to 25%, and a ratio of 1:3 retained 75% of the selenocystine-alone signal. The proportional change in fluorescent signal with varying To further assess the transport selectivity of SLC3A1/SLC7A9 transporters, we utilized known basic amino acids substrates, such as lysine, arginine, and ornithine, to compete with selenocystine. As depicted in Fig. 3B, a combination of 1/2 ornithine and 1/2 selenocystine blocked over half of the transport signal, whereas a mixture of 3/4 ornithine and 1/4 selenocystine obstructed 90% of the signal, indicating that ornithine has a higher selectivity. Similarly, Fig. 3C and 3D illustrate that the selectivity for arginine and lysine is significantly greater than for selenocystine. Notably, as shown in Fig. 3E and 3F, neutral amino acids such as glycine, alanine, and serine, as well as acidic amino acids such as aspartic acid and glutamic acid, did not inhibit selenocystine transport. This hierarchy of substrate preference aligns with previous studies based on isotopes (12–14), which highlights the reliability of the selenocystine-based fluorescence assay in evaluating transport selectivity.

**Figure 3.**
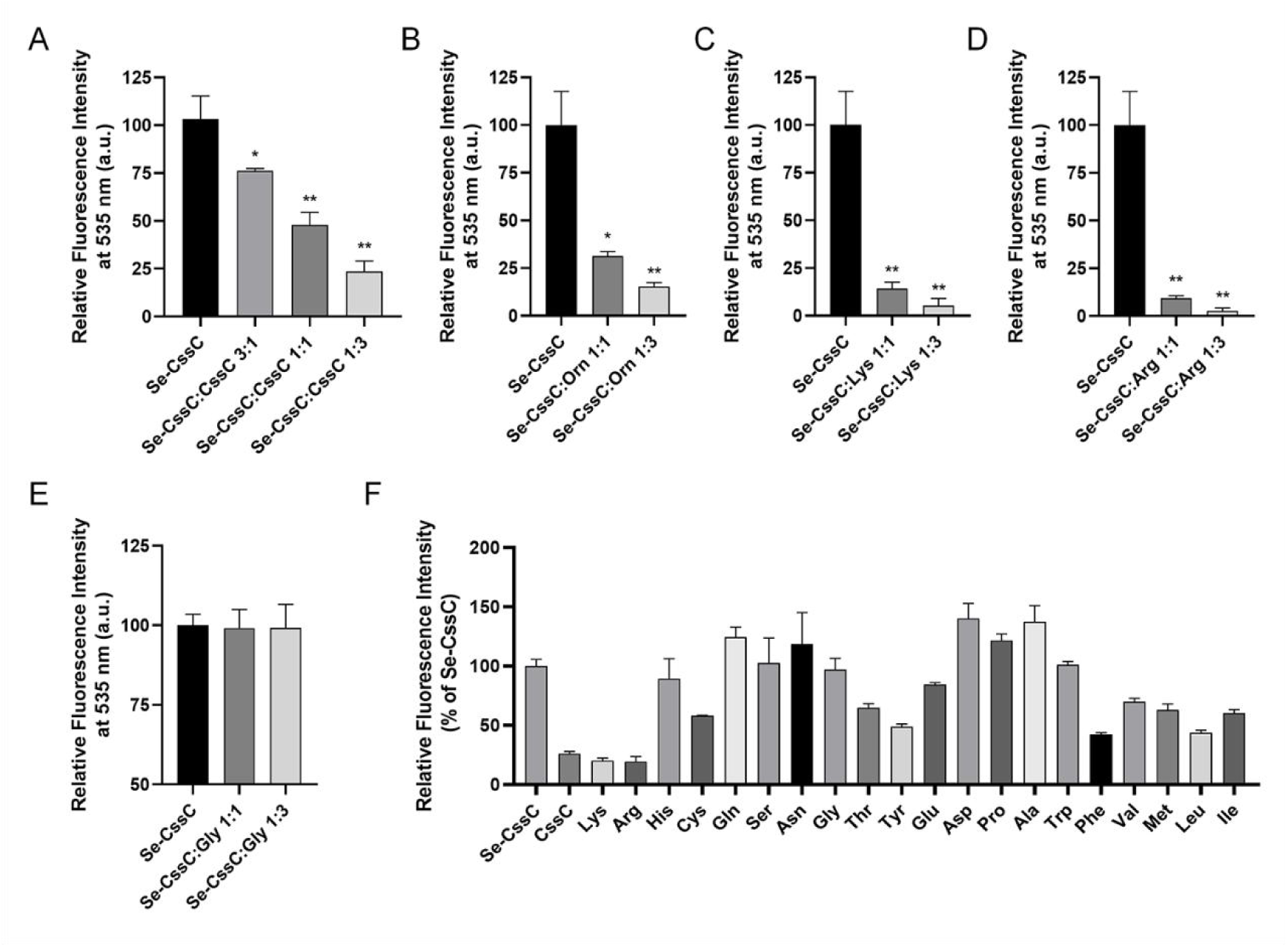
Transport function and substrate selectivity of SLC3A1/SLC7A9. (A-E) Competitive inhibition of selenocystine transport by unlabeled amino acids. SLC3A1/SLC7A9-expressing HEK293 cells were treated with selenocystine and competing amino acids (total concentration: 400 μM) at ratios of 4:0 (control), 2:2, or 1:3 (selenocystine: competitor). Transport activity is expressed as a percentage relative to the control condition (selenocystine only, 4:0 ratio). (A) Competition with cystine. (B) Competition with ornithine. (C) Competition with lysine. (D) Competition with arginine. (E) Competition with glycine. (F) Inhibition of selenocystine transport (200 μM initial extracellular concentration) in competition assays containing 1 mM unlabeled amino acid (as indicated, with the exception of tyrosine, which was used at 600 μM because of low solubility). Transport is displayed as percentage of no competing amino acid (only selenocystine). \**P* < 0.05; ** *P*<0.01.

To further validate the assay, we examined several clinically relevant missense variants of SLC7A9 and SLC3A1 previously characterized using radioisotope-based methods. Specifically, we selected variants representative in terms of pathogenic classification and population frequency based on gnomAD and ClinVar databases.

SLC7A9 variant A182T (rs79389353), classified as having “Conflicting classifications of pathogenicity” and occurring at the highest frequency (0.004069) within this classification, along with A70V (rs769448665), classified as “Likely pathogenic” with a gnomAD frequency of 0.000029 (also the highest within this classification), have both been associated with mild functional impairment (5, 15). Consistent with previous reports, both A182T and A70V demonstrated similarly mild impairments in the Selenocystine-Based Fluorescence Assay (Fig. 4A and 4B). Variants SLC7A9 G105R (rs121908480) and R333W (rs121908484), classified as “Pathogenic/Likely pathogenic” with respective gnomAD frequencies of 0.000492 and 0.000208 (the two highest frequencies within their classification), were previously associated with moderate to severe functional impairment (5, 7). Our assay results corroborated these findings, clearly showing moderate to severe transporter dysfunction (Fig. 4C and 4D). Further, we evaluated SLC7A9 P482L (rs146815072) classified as “Pathogenic” with a frequency of 8.98505E-05 (highest within this classification), and SLC3A1 M467T (rs121912691) classified as “Pathogenic/Likely pathogenic” with a frequency of 0.003161 (highest within this classification). Additionally, two other common severe loss-of-function variants, SLC7A9 V170M (rs121908479, frequency 4.95677E-06) and A354T (rs939028046, frequency 3.22429E-05), were analyzed. All four variants resulted in near-complete loss of transport function, aligning well with previously published functional data (5, 7, 15–17). Consistent with these expectations, the Selenocystine-Based Fluorescence Assay revealed a significant reduction or complete absence of transport activity for these variants (Fig. 4E and 4F).

**Figure 4.**
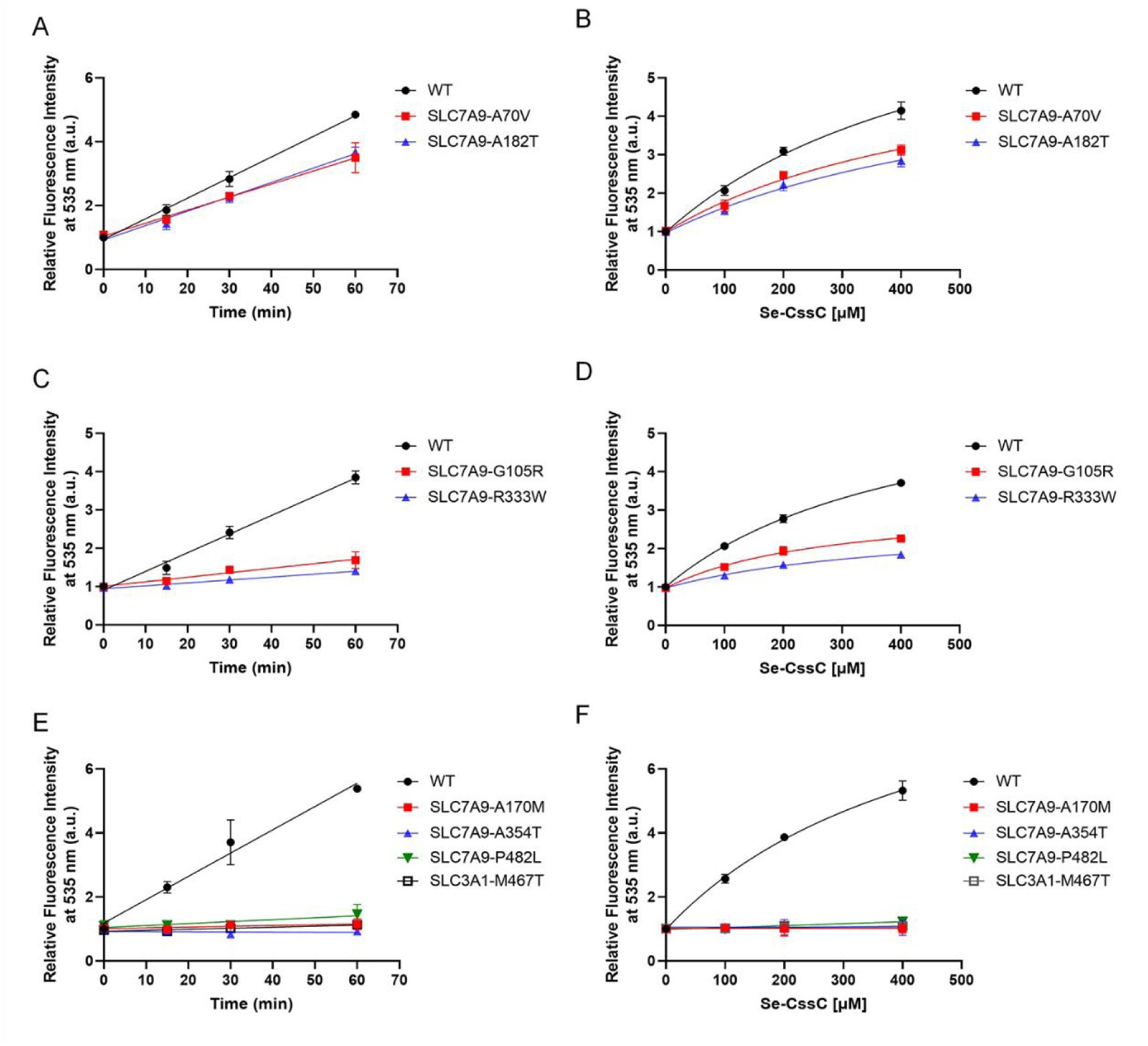
Selenocystine Transport Capacity of Wild-Type and Clinically Reported Mutants of SLC7A9 and SLC3A1. (A-F) HEK293 cells co-expressing wild-type SLC3A1 with either wild-type SLC7A9 or indicated mutants were treated with selenocystine. Transport activity (fluorescence intensity) is expressed as fold change relative to basal (untreated control). (A) Time-dependent uptake of 200 μM selenocystine by wild-type SLC7A9, SLC7A9-A70V, or SLC7A9-A182T. (B) Dose-dependent uptake of selenocystine (0–400 μM, 30 min incubation) for wild-type SLC7A9, SLC7A9-A70V, or SLC7A9-A182T. (C–D) Time-dependent (C) and dose-dependent (D) uptake assays for wild-type SLC7A9, SLC7A9-G105R, or SLC7A9-R333W. (E–F) Time-dependent (E) and dose-dependent (F) uptake assays for wild-type SLC3A1/SLC7A9, SLC7A9-A170M, SLC7A9-R333W, SLC7A9-P482L, or SLC3A1-M467T.

Collectively, these results confirm that the Selenocystine-Based Fluorescence Assay accurately and reliably captures the spectrum of transporter dysfunction caused by clinically significant mutations in SLC7A9 and SLC3A1, underlining its robustness and utility in functional diagnostics.

### 3.3 SLC7A13 (AGT1) has no ability to transport selenocystine

An unsolved paradox is that SLC3A1 is highly expressed in the S3 segment, the renal late proximal tubules, whereas SLC7A9 expression is highest in the S1 segment. Researchers came up a hypothesis that there are missing transporter partners for SLC3A1 and/or SLC7A9. SLC7A13 (AGT1) was proposed to be the missing cystine transporter partner for SLC3A1, as AGT1 were colocalized with SLC3A1 in the S3 segment and purified AGT1-SLC3A1 heterodimer were reported to transports cystine, aspartate, and glutamate(18). Controversially, the proposal that SLC7A13 serves as the missing partner for SLC3A1 has faced scrutiny due to conflicting evidence(19). We therefore used our assay to reexamine the ability of SLC7A13 on Selenocystine transporting. As the results shown in Fig. 5A and 5B, our evidence supports SLC7A13 is not a selenocystine/cystine transporter. As a control, the SLC7A11 combined SLC3A2 not SLC3A1 induced Selenocystine transporting.

**Figure 5.**
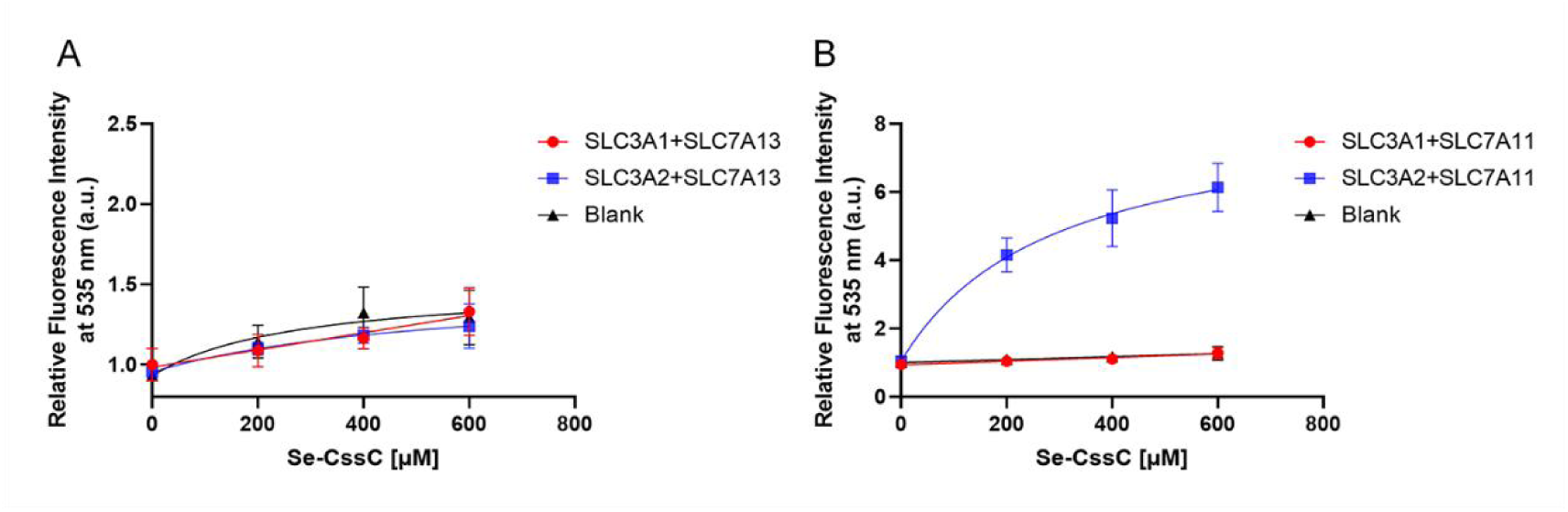
Selenocystine Transport activity of other putative transporter complexes. HEK293 cells transfected with SLC3A1/SLC7A13 (A), SLC3A2/SLC7A13 (A), SLC3A1/SLC7A11 (B), or SLC3A2/SLC7A11 (B) were incubated with increasing concentrations of selenocystine for 30 minutes. Untransfected cells (blank) served as controls. Fluorescence intensity (reflecting selenocystine uptake) was measured using a multifunctional microplate reader. Data are expressed as fold change relative to untransfected controls.

### 3.4 Rapidly identify AlphaFold-predicted structural changes that induce impairment

Recently, AlphaFold2/3 have shown remarkable accuracy in predicting protein structures(20, 21). These computational structure prediction tools, when combined with small molecular docking tools, provides the foundation for a mechanistic understanding of cystinuria-related mutations. To assess the effectiveness of these predictions in identifying transport impairments through disrupting substrate binding or transporter channel, we compared the predicted structures of wild-type and mutant SLC7A9 variants. As illustrated in Fig. 6A, the SLC7A9 P482L mutation results in a bulky side chain that obstructs the cystine transport pathway, while P482G does not. Consistent with the functional assay results reported by Y Shigeta et al.(16), our fluorescence assay (Fig. 6B) indicates that the P482L mutation retains only 5% activity compared to the wild-type (WT) complex. In contrast, P482G maintains full capacity at 100%. These outcomes elucidate the mechanism behind the loss of function in the P482L mutant, a phenomenon that was previously unresolved (16). The SLC7A9 mutant A354T exhibited negligible transport activity (Fig.4E and 4F). It disrupted TM rotations and caused R326 to protrude from the side of the transport channel, thereby impeding the channel’s ability to transport amino acids (Fig. 6C). These findings highlight the potential of integrating structural predictions with functional assays to rapidly and accurately identify mutations that impair cystine transport.

**Figure 6.**
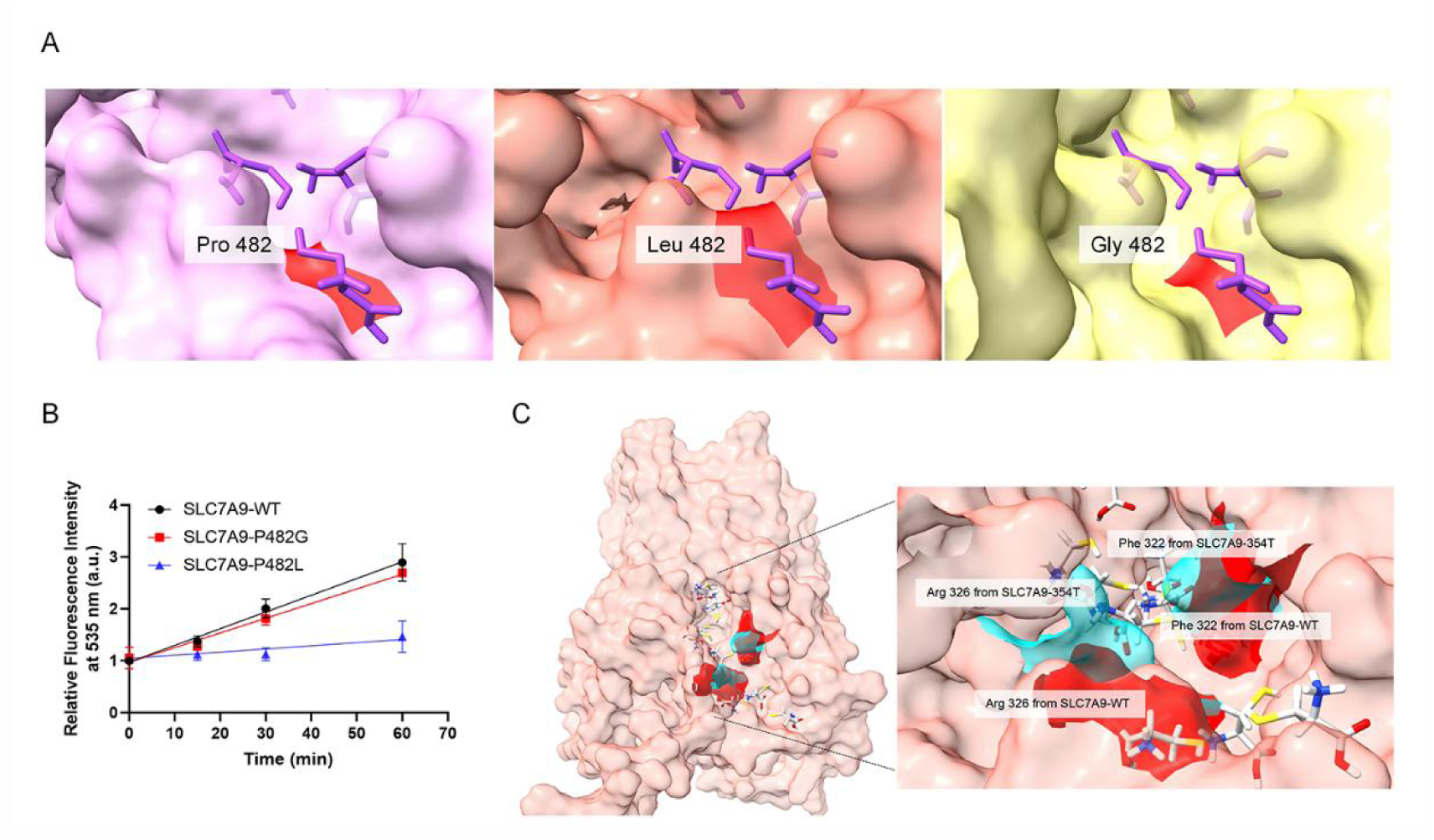
Structure prediction and functional identification of mutant of SLC7A9 by selenocystine uptake assay. (A) Surface structural comparison of wild-type and mutant b0,+AT at position 482. Left to right: Surface representations of wild-type b0,+AT, P482L mutant, and P482G mutant. Residue 482 (red) highlights conformational changes induced by proline-to-leucine (P482L) or proline-to-glycine (P482G) substitutions. (B) HEK293 cells co-expressing wild-type SLC3A1 with either wild-type SLC7A9 or the indicated mutants were treated with 200 μM selenocystine for the indicated duration. Transport activity (fluorescence intensity) is expressed as fold change relative to basal (untreated control). (C) Structural perturbation of residues 322 and 326 in the A354T mutant of b0,+AT. Surface representation of wild-type b0,+AT (background, red) with residues 322 (right, red) and 326 (left, red). The A354T mutant induces structural rearrangements at these positions, shown in cyan. The mutation at position 354 (not displayed) disrupts local interactions, altering the conformation of distal residues 322 and 326.

## 4. Discussion

Traditional approaches for evaluating cystine transporter functionality, such as radioisotope-labeled amino acid uptake assays and mass spectrometry-based cystine quantification, present significant limitations in a clinical context. While over 278 and 182 pathogenic mutations have been reported in the SLC3A1 and SLC7A9 genes respectively, only a small fraction have been functionally characterized to date (Data from HGMD, January 2025 version)(22). This gap is largely due to the impracticality of conventional assays for routine use, radioisotope methods carry safety risks and require specialized facilities and mass spectrometry, demands expensive instrumentation and technical expertise not readily available in all laboratories (1). Our study introduces a novel fluorescence-based uptake assay utilizing selenocystine as a cystine analog, enabling sensitive, cost-effective, and scalable assessment of SLC3A1/SLC7A9 transporter function. This approach eliminates the need for radioactive substrates and capital-intensive equipment, relying instead on standard microplate readers and non-hazardous reagents. Notably, fluorometric detection methods are generally regarded as simple, rapid, and highly sensitive, which is consistent with our findings(23). In agreement with prior studies on the cystine/glutamate antiporter(11), we confirmed that selenocystine closely mimics cystine in transport assays, supporting the validity of our fluorescence-based method for detecting cystine transport activity.

A major finding of this study is the ability of our fluorescence assay to evaluate the functional impact of clinically relevant variants in cystinuria. Specifically, variants previously associated with mild cystinuria phenotypes (e.g. SLC7A9 A70V and A182T) showed minimal transport impairment in our assay, whereas intermediate variants like SLC7A9 G105R and R333W led to moderate reductions in transport efficiency. Severe loss-of-function mutations including SLC7A9 P482L, V170M and A354T, and SLC3A1 M467T, exhibited near-complete loss of transport activity, aligning with patients’ more severe clinical phenotype (5, 24).

These results not only validate our assay as a reliable tool for assessing transporter activity, but also demonstrate its capacity to stratify variants by functional severity. Such stratification could be clinically significant: the ability to differentiate mild, moderate, and severe functional defects suggests that the assay may help predict disease severity and stone risk in patients based on their specific genotype. Indeed, our data resolve a longstanding controversy by showing that SLC7A13 (formerly AGT1) does not transport cystine, clarifying that only the SLC3A1/SLC7A9 complex, and not alternative transporters, is relevant in cystinuria (18, 19). Our experimental results indicate that SLC7A13 lacks cystine transport capacity, contradicting prior hypotheses. This clarification underlines the assay’s utility in pinpointing the true molecular drivers of the disease.

In addition to functional measurements, integrating structural predictions provided mechanistic insight into how particular mutations cause transporter dysfunction. AlphaFold-based models helped explain, for example, how the SLC7A9 P482L mutation introduces a steric block in the transport pathway, correlating with the markedly reduced cystine uptake observed experimentally. Similarly, the A354T variant was found to disrupt transmembrane domain rotations, leading to transporter inactivation. These findings underline the potential of combining structural modeling with functional assays for rapid and accurate characterization of cystine transport impairments.

In summary, our fluorescence-based selenocystine transport assay combining structural predictions provides a sensitive and reliable tool for assessing cystine transporter function and identifying transport-deficient variants. Future studies should focus on validating its clinical applicability and integrating it into high-throughput screening platforms to facilitate cystinuria research and therapeutic development.

## Data availability statement

The data that support the findings of this study are available upon reasonable request from the corresponding author.

## Author Contribution Statement

X.H. took the responsibility for Funding acquisition, Investigation. X.Q., X.Z., H.Z., J.S., and R.Q. participated in the investigation. X.C. provided Resource. J.L. and L.Y. provided guidance for thesis and financial support. L.C.: as the corresponding author, took charge of the Conceptualization; Supervision; Writing-review & editing. All authors read and approved the final manuscript.

## Research ethics

Not applicable.

## Informed consent

Not applicable.

## Acknowledgements

The authors acknowledge the support from the Scientific Research Center, Hangzhou Medical College.

